# Mindcontrol: A Web Application for Brain Segmentation Quality Control

**DOI:** 10.1101/090431

**Authors:** Anisha Keshavan, Esha Datta, Ian McDonough, Christopher R. Madan, Kesshi Jordan, Roland G. Henry

**Affiliations:** Department of Neurology, University California, San Francisco, USA; UC Berkeley - UCSF Graduate Program in Bioengineering, San Francisco, USA; The University of Alabama, Tuscaloosa, AL, United States of America; Department of Psychology, Boston College, Chestnut Hill, MA, USA; Department of Radiology and Biomedical Imaging, University California, San Francisco, CA, USA

## Abstract

Tissue classification plays a crucial role in the investigation of normal neural development, brain-behavior relationships, and the disease mechanisms of many psychiatric and neurological illnesses. Ensuring the accuracy of tissue classification is important for quality research and, in particular, the translation of imaging biomarkers to clinical practice. Assessment with the human eye is vital to correct various errors inherent to all currently available segmentation algorithms. Manual quality assurance becomes methodologically difficult at a large scale - a problem of increasing importance as the number of data sets is on the rise. To make this process more efficient, we have developed Mindcontrol, an open-source web application for the collaborative quality control of neuroimaging processing outputs. The Mindcontrol platform consists of a dashboard to organize data, descriptive visualizations to explore the data, an imaging viewer, and an in-browser annotation and editing toolbox for data curation and quality control. Mindcontrol is flexible and can be configured for the outputs of any software package in any data organization structure. Example configurations for three large, open-source datasets are presented: the 1000 Functional Connectomes Project (FCP), the Consortium for Reliability and Reproducibility (CoRR), and the Autism Brain Imaging Data Exchange (ABIDE) Collection. These demo applications link descriptive quality control metrics, regional brain volumes, and thickness scalars to a 3D imaging viewer and editing module, resulting in an easy-to-implement quality control protocol that can be scaled for any size and complexity of study.

## 1. Background

Imaging biomarkers derived from MRI play a crucial role in the fields of neuroscience, neurology, and psychiatry. Estimates of regional brain volumes and shape features can track the disease progression of neurological and psychiatric diseases such as Alzheimer’s disease [1, 2], Parkinson’s disease [3], schizophrenia [4], depression [5], autism [6], and multiple sclerosis [7]. Given recent increases in data collection to accommodate modern precision-medicine approaches, assuring the quality of these biomarkers is vital as we scale their production.

Various semi-automated programs have been developed to estimate MRI biomarkers. While these applications are efficient, errors in regional segmentation are inevitable, given several methodological challenges inherent to both technological and clinical implementation limitations. First, the quality of the MRI scan itself due to motion artifacts or scanner instabilities could blur and distort anatomical boundaries [8, 9, 10, 11]. Differences in MRI hardware, software, and acquisition sequences also contribute to contrast differences and gradient distortions that affect tissue classification, which makes combining datasets across sites challenging [12]. An additional source of error comes from parameter selection for segmentation algorithms; different parameter choices can translate to widely varying results [13]. Furthermore, many MR segmentation algorithms were developed and tested on healthy adult brains; applying these algorithms to brain images of children, the elderly, or those with pathology may violate certain assumptions of the algorithm, resulting in drastically different results.

Several quality assurance strategies exist to address segmentation errors. In one approach, researchers flag low-quality scans prior to analysis by viewing the data before input to tissue classification algorithms. However, identifying “bad” datasets using the raw data is not always straightforward, and can be prohibitively time consuming for large datasets. Pre-processing protocols have been developed to extract metrics that can be viewed as a cohort-level summary from which outliers are selected for manual quality-assurance. For example, by running the Preprocessed-Connectomes Project’s Quality Assurance Protocol (PCP-QAP) [14], researchers can view summary statistics that describe the quality of the raw data going into the algorithm and automatically remove subpar images. However, these metrics are limited because segmentation may still fail even if the quality of the scan is good. Another quality assurance strategy is to plot distributions of the segmentation output metrics themselves and remove any outlier volumes. However, without manual inspection, normal brains that naturally have very small or large estimates of brain size or pathological brains with valid segmentations may be inappropriately removed. Ideally, a link would exist between scalar summary statistics and 3D/4D volumes. Such a link would enable researchers to prioritize images for labor-intensive quality control (QC) procedures; to collaborate and organize QC procedures; and to understand how scalar quality metrics, such as signal to noise ratio, relate to the actual image and segmentation. In this report, we present a collaborative and efficient MRI QC platform that links group-level descriptive statistics to individual volume views of MRI images.

We propose an open source web-based brain quality control application called Mindcontrol: a dashboard to organize, QC, annotate, edit, and collaborate on neuroimaging processing. Mindcontrol provides an intuitive interface for examining distributions of descriptive measures from neuroimaging pipelines (e.g., surface area of right insula), and viewing the results of segmentation analyses using the Papaya.js volume viewer (https://github.com/rii-mango/Papaya). Users are able to annotate points and curves on the volume, edit voxels, and assign tasks to other users (e.g., to manually correct the segmentation of a particular image). The platform is pipeline agnostic, meaning that it can be configured to quality control any set of 3D volumes regardless of what neuroimaging software package produced it. In the following sections, we describe the implementation details of Mindcontrol, as well as its configuration for three open-source datasets, each with a different type of neuroimaging pipeline output.

## 2. Software Design and Implementation

### 2.1 Design Principles

Mindcontrol was developed with several design requirements. Mindcontrol must be easily accessible from any device, such as a Mac, Windows or even a tablet. Therefore, the best option was to develop a web application. Most tablets have limited storage capacity, so space-minimizing specifications were established. A dependence on cloud-based data storage was specified to accommodate large neuroimaging datasets without needing local storage. To efficiently store annotations and edited voxels, Mindcontrol only stores the changes to files, rather than whole-file information, on its database. Researchers must be able to QC outputs from any type of neuroimaging software package, so Mindcontrol was specified to flexibly accommodate any file organization structure, with configurable “modules” that can contain any type of descriptive statistics and 3D images. Mindcontrol configuration and database updates must require minimal Javascript knowledge, since Matlab/Octave, Python, R, and C are primarily used in the neuroimaging community for data analysis. Finally, changes to the database(like the addition of new images), changes in descriptive measures, and new edits/annotations, should be reflected in the application in real-time to foster collaboration.

### 2.2 Server Back-End Framework

Mindcontrol is built with Meteor (http://www.meteor.com), a full-stack javascript web-development platform. Meteor features a build tool, a package manager, the convenience of a single language (javascript) to develop both the front-and back-end of the application, and an abstracted implementation of full-stack reactivity. Data is transferred “over the wire” and rendered by the client (as opposed to the server sending HTML), which means that changes to the database automatically trigger changes to the application view. For example, as soon as a user finishes implementing QC procedures on an image and clicks “save”, all other users can see the changes. A diagram of this process is provided in Figure 1.

**Figure 1:**
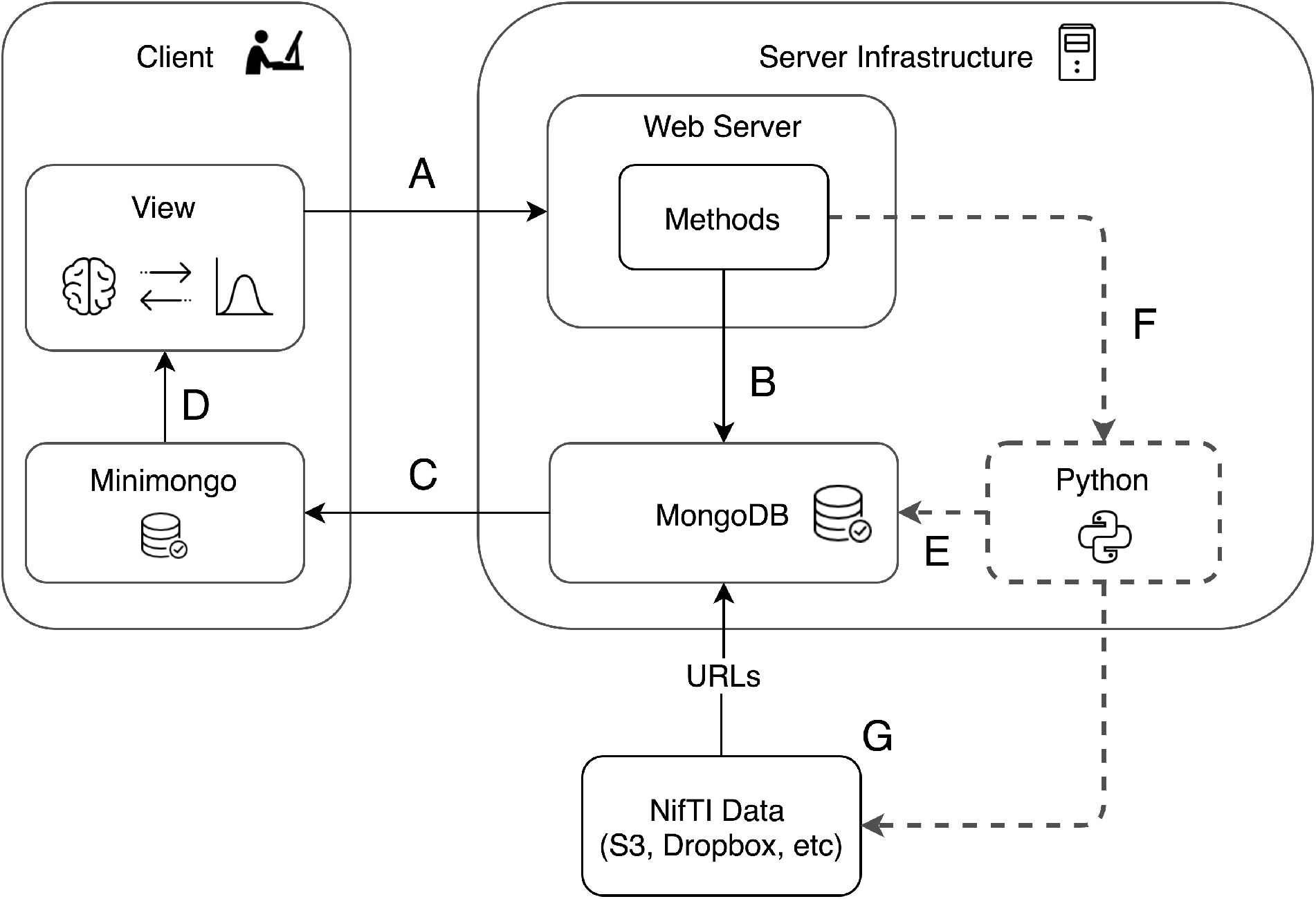
This diagram shows the different components of the Mindcontrol application. A) The client sends information, such as annotations and edits, to the server. B) The server calls a method that updates the mongoDB backend. C) When the back-end MongoDB database changes, these changes are automatically pushed to the minimongo database on the client. D) A change to the minimongo database automatically re-renders the view on the client. E) Users can optionally push changes to the client view via the MongoDB with Python MongoDB drivers. Drivers for C, C++, Scala, Java, PHP, Ruby, Perl, and Node.js are also available through MongoDB. F) Developers can optionally write server methods to launch Python or command-line processes that, in turn, use user annotations and edits to re-process images and update the MongoDB with new results. G) Imaging data (in NifTI format) is stored on an external server, such as Amazon S3 or Dropbox, and URLs to the images are stored in the MongoDB.

### 2.3 Client-Side Features

The user interface consists of a dashboard view and an imaging view, as shown in Figures 2 and 3, respectively. The primary dashboard view consists of processing module sections, a query controller, data tables, and descriptive statistic visualizations. Each entry in the table is a link that, when clicked, filters all tables on the page. The filters or queries can be saved, edited, and loaded in the query controller section, as shown in Figure 4.

**Figure 2:**
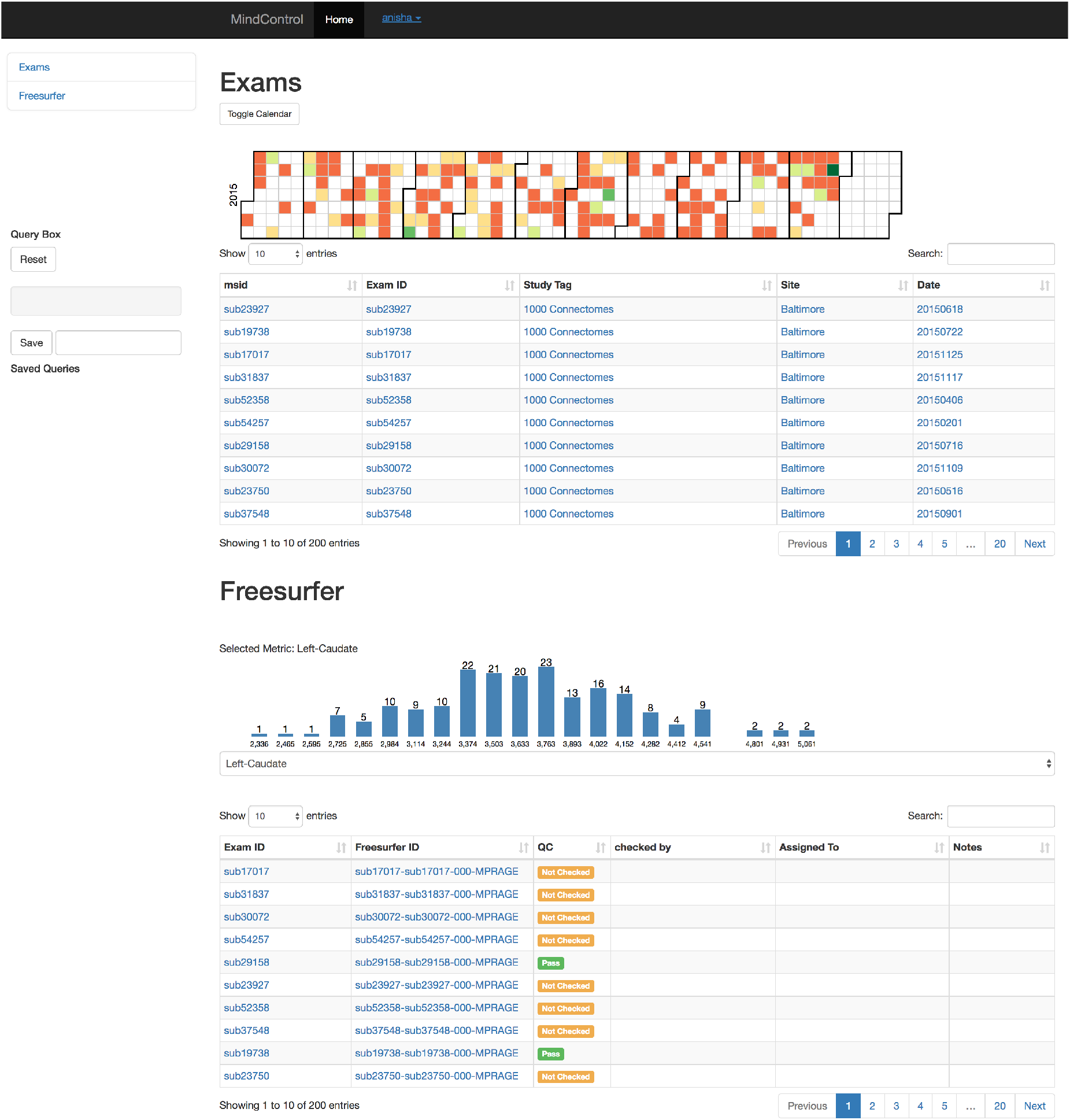
This figure shows the Mindcontrol layout configured to quality check Freesurfer outputs from the 1000 Functional Connectomes Project (FCP). Part A shows the module navigator, which links to the different processing modules on the dashboard. Part B shows the different exams and the dates they were acquired as a heatmap, where green is more and orange is less scans collected on a given day. (For demonstration purposes, the dates depicted here do not reflect the actual dates the data were collected for the FCP, since this information was not provided at the time.) Clicking on data in any column of the exam table filters the data by that column. For example, clicking the site “Milwaukee” reduces both the “Exams” and the “FreeSurfer” tables to only show subjects from Milwaukee. Part C shows the Freesurfer table and regional volume distribution of the left caudate. A drop-down menu allows users to switch the descriptive metric. Clicking on a value in FreeSurfer ID column brings the user to the imaging view, as shown in Figure 3, where users can evaluate and annotate the quality status of the image. The value of the label in the “QC” column changes instantaneously due to Meteor’s built in full-stack reactivity.

**Figure 3:**
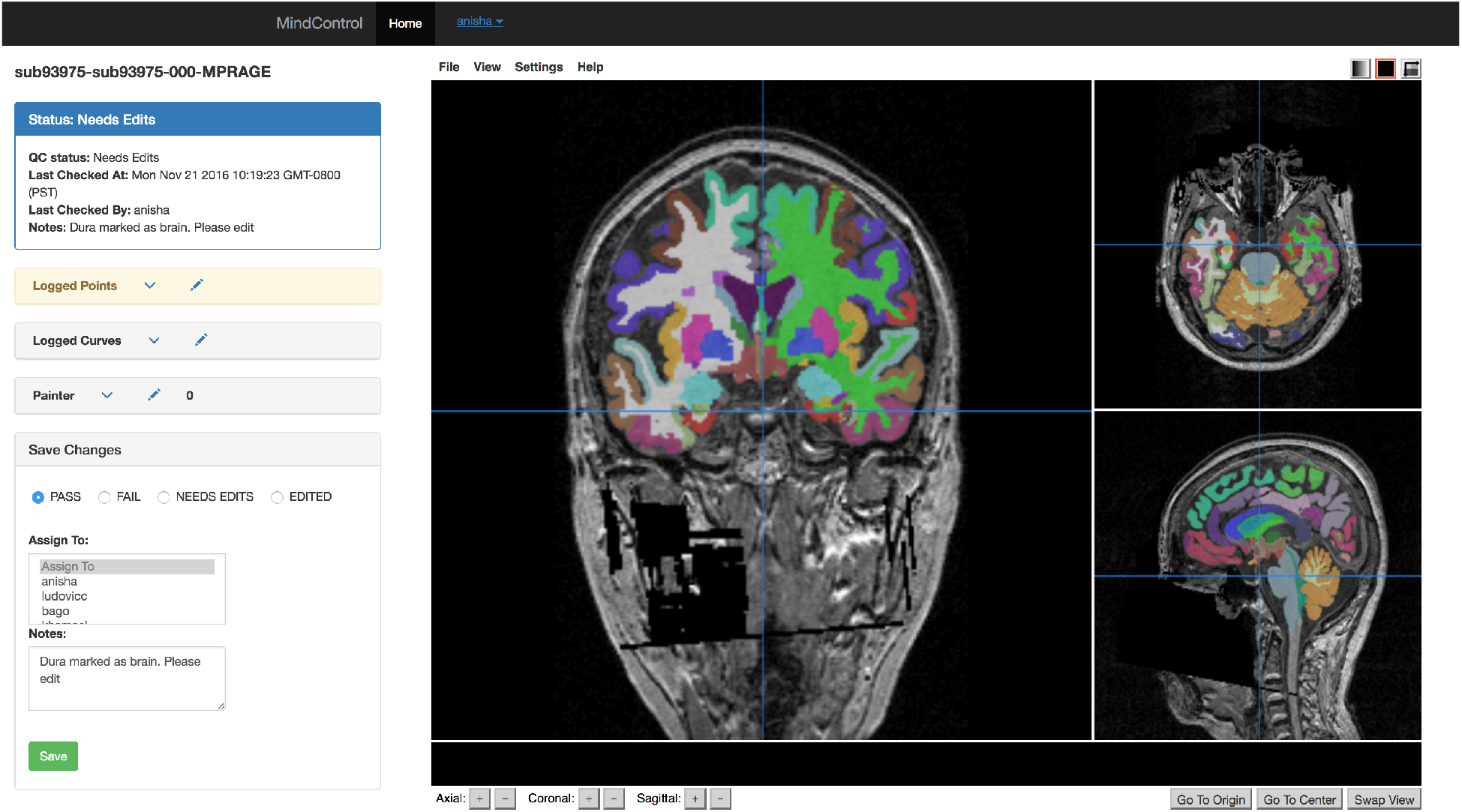
The imaging view of mindcontrol consists of a panel on the left-hand side that contains the QC status; a point annotation menu; a curve annotation menu; a voxel editing menu; and an editable sub-panel for QC status, notes, and editor assignment. On the right-hand side, the base MRI anatomical MPRAGE image is displayed with an overlay of the Freesurfer segmentation outputs using the Papaya.js viewer.

**Figure 4:**
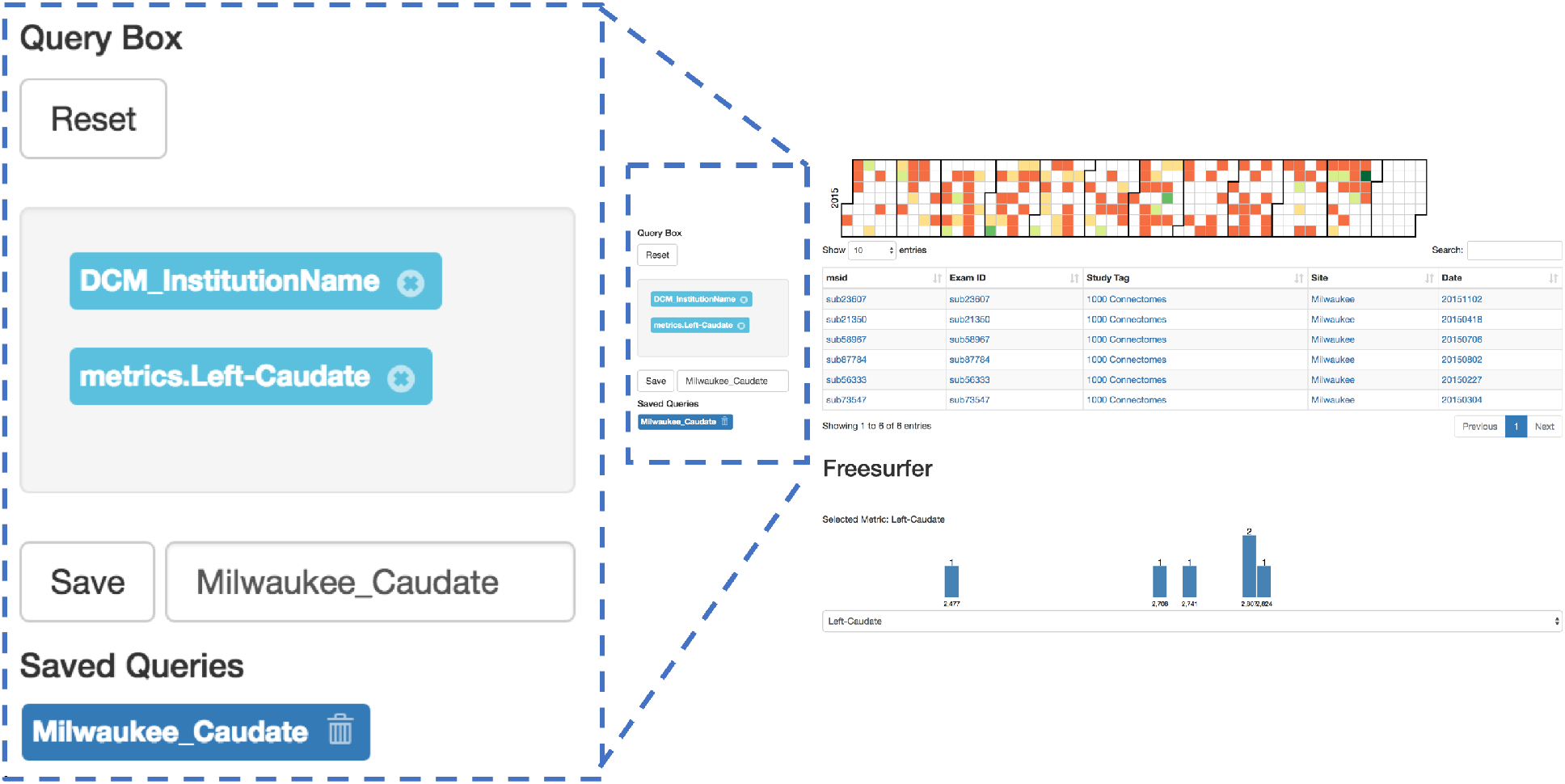
The query controller shows the different filters that have been applied to this dataset. In this example, the exams have been filtered by institution (“Milwaukee”) and by a range of left caudate volumes (brushed from the histogram). Clicking the “x” next to the filter removes it, and the view updates. Queries can be saved and reloaded by providing a name in the text-entry box and clicking “Save”. “Reset” removes all filters to show the whole dataset.

Descriptive statistics are visualized using the D3 library (https://d3js.org/). Currently, two visualizations are provided: a calendar view of a heatmap that shows a histogram of the number of exams collected on a given day and 1D histograms of scalar metrics with dimensions that are swappable using a dropdown menu, as shown in Figure 2. Both histogram plots interactively filter the data tables below. Clicking on a particular date on the date-histogram plot filters all tables by the exams collected on that particular date. Users are able to “brush” sections of the 1D histogram to filter all tables with exams that meet requirements of values within that range (see Figure 5).

**Figure 5:**
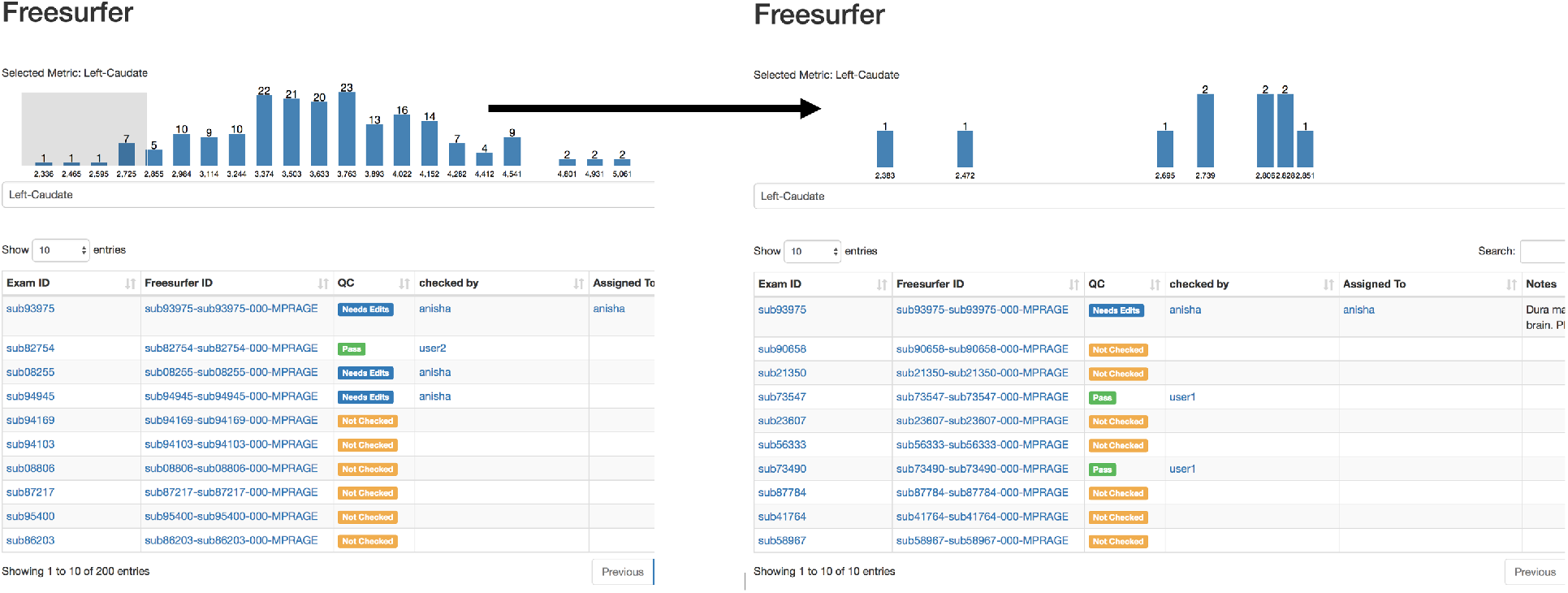
This demonstrates the interactive brushing feature of Mindcontrol histograms. On the left, the user has brushed the tail end of the left caudate volume distribution from Freesurfer. On the right, the histogram has been redrawn with data from the brushed range, and the table beneath filtered from 200 entries to 14 entries based on the brushed caudate volumes.

The imaging view is shown in Figure 3. The left-side column includes a section to label an image as “Pass”, “Fail”, “Edited”, or “Needs Edits” and to provide notes. The status bar at the top-left portion updates instantaneously with information on which user checked the image, the quality status of the image, and when it was last checked. Users are also able to assign edits to be performed by other users on the system; for example, a research assistant can perform a general QC and assign difficult cases to a neuroradiologist. On the right-hand side, the Papaya.js viewer (http://rii-mango.github.io/Papaya/) is used to display the NifTI volumes of the original data and FreeSurfer segmentations.

Annotations of points and curves are shown in Figure 6. Using the *shift* key, users can click on the image to annotate points or select the “Logged Curves” toolbar. By *shift+click and dragging*, users can draw curves. Keyboard and mouse shortcuts provided by the Papaya.js viewer, along with Mindcontrol, include toggling overlays (*zz*) and undoing annotations (*dd*). Figure 7 shows the editing (“Painter”) panel of the imaging view. Users set paintbrush values and *shift+click and drag* to change voxel values. For point and curve annotations and voxel editing, the images themselves are not changed, but world x,y,z coordinates, along with annotation text or paintbrush values, are saved to the mongo database when the user clicks “save”. Custom offline functions may be written to apply editing to images: for example, to implement pial surface edits from FreeSurfer.

**Figure 6:**
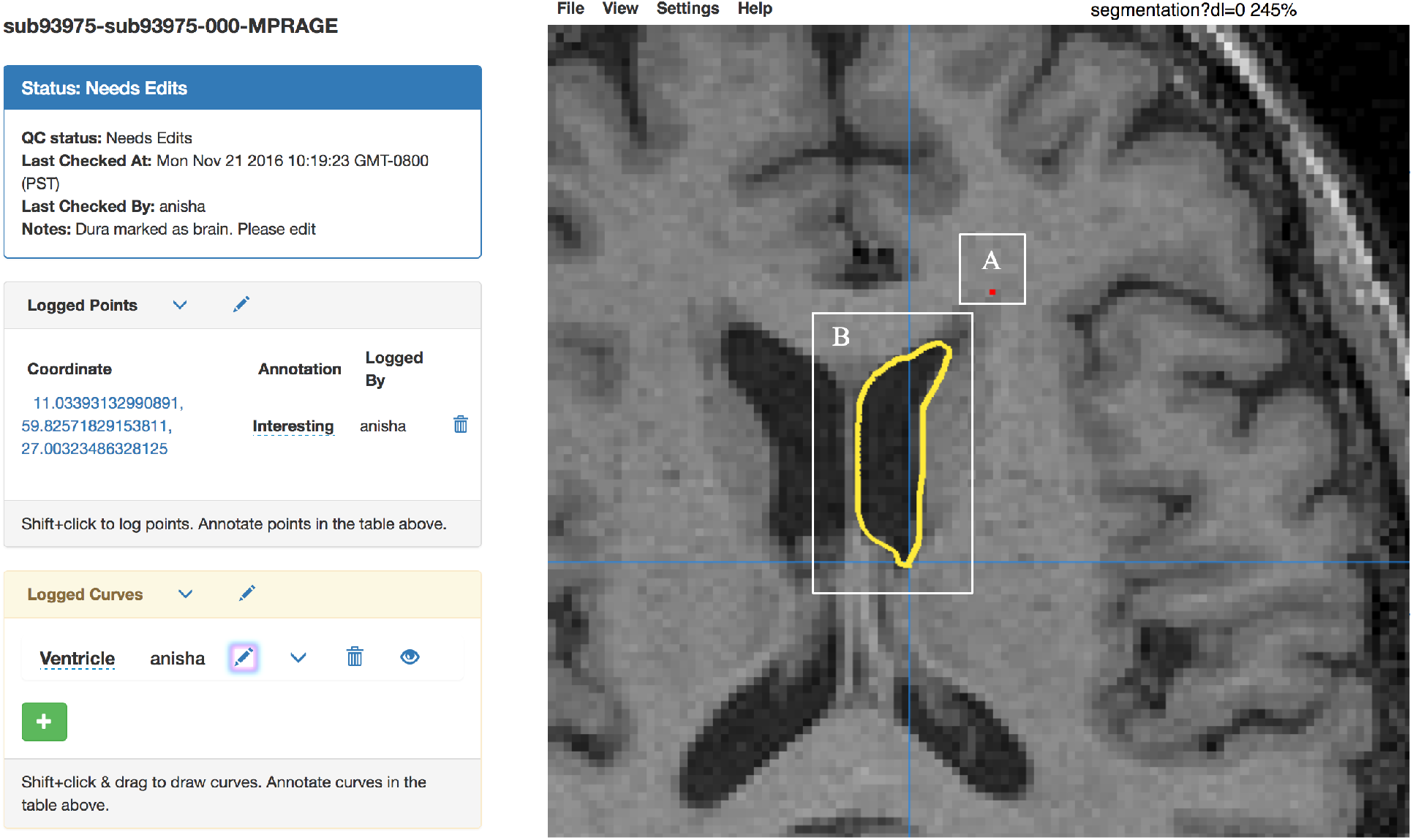
The annotations panel can be used to annotate a single point (shown in red, part A) and curves (shown in B). When annotating points, the user is shown the selected x,y,z world coordinates and is able to name the annotation. In the curve annotation panel on the left sidebar, the user is able to name the curve and add/remove curves. Keyboard shortcuts: “dd” removes the previous annotation and “zz” toggles the segmentation overlay.

**Figure 7:**
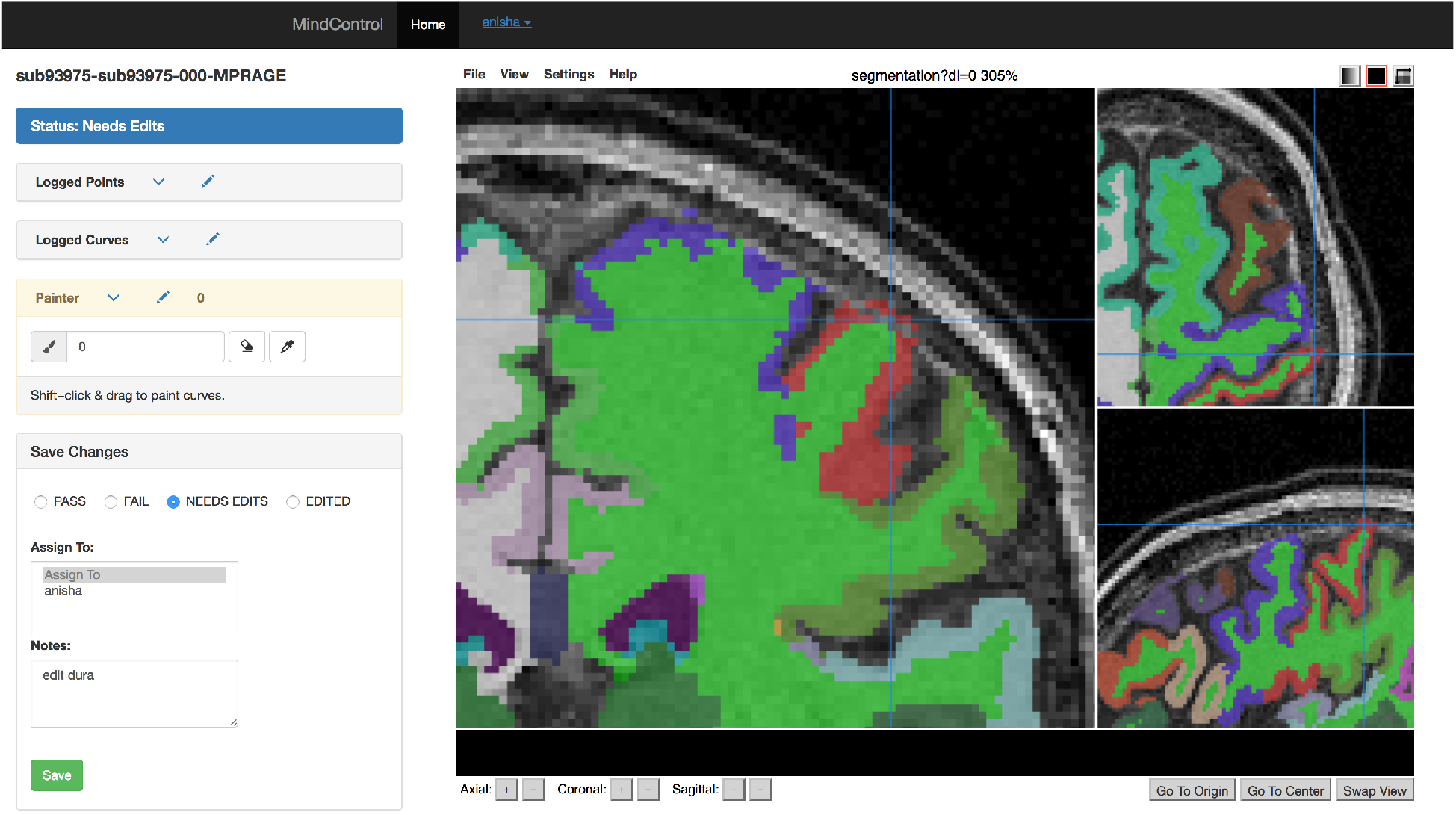
The editing panel on the left shows the “Painter” toolbox in yellow, where users can input brush values or use the eyedropper tool to set the value to that of a clicked label. The eraser icon sets the brush value to 0, to delete or erase voxels. In the image above, the Freesurfer segmentation is being edited by erasing the voxels missclassified as dura.

### 2.4 Configuration Details

Mindcontrol can be configured for a study’s specific needs be specifying a configuration JSON file. The configuration file describes processing modules by module names, the columns to display in the data table below, and the type of graph to display (Date histogram or 1D histogram). Images must be hosted on a separate server or a content delivery network (CDN) and the Mindcontrol database populated with URLs to these images. An initial JSON file can be specified to populate the Mindcontrol MongoDB with entries on startup if the database is empty. Instructions and example JSON schema for the configuration file and the database entries can be found at https://github.com/akeshavan/mindcontrol/wiki along with a Python function to access the MongoDB, which can be used to write custom editing scripts and externally update the database.

## 3. Examples/Applications

Mindcontrol configurations were developed for selected data from the 1000 Functional Connectomes project (FCP), the consortium for reliability and reproducibility (CoRR), and the Autism Brain Imaging Data Exchange (ABIDE) Collection I.

The FCP consists of 1414 resting state fMRI and corresponding structural datasets collected from 35 sites around the world [15], which have been openly shared with the public. The purpose of the FCP collaboration is to comprehensively map the functional human connectome, to understand genetic influences on brain’s structure and function, and to understand how brain structure and function relate to human behavior [15]. Segmentation of 200 selected FCP anatomical images from Baltimore, Bangor, Berlin, ICBM, and Milwaukee was performed with Freesurfer (recon-all) version 5.3.0 [16] using the RedHat 7 operating system on IEEE 754 compliant hardware at UCSF. Regional volumes of subcortical and cerebellar regions were computed. Cortical volumes, areas, and thicknesses were also computed and averaged across hemispheres. Scan dates were simulated in order to demonstrate the date histogram shown in Figure 1B. The original anonymized T1-weighted images, along with the aparc+aseg output from Freesurfer, were converted to the compressed NifTI (.nii.gz) format and uploaded to Dropbox for the purpose of visualization within Mindcontrol. The Mindcontrol database was populated with URLs to these images, along with their corresponding FreeSurfer segmentation metrics. The demo of the FCP data is located at https://github.com/akeshavan/mindcontrol/wiki.

More recently, researchers have developed the Preprocessed-Connectomes Project’s Quality Assurance Protocol (PCP-QAP) software, to provide anatomical and functional data quality measures in order to detect low-quality images before data processing and analysis [14]. Some metrics include contrast-to-noise ratio, signal-to-noise ratio, voxel smoothness, percentage of artifact voxels, foreground-to-background energy ratio, and entropy focus criterion [14]. The PCP-QAP protocol has been run on the Consortium for Reliability and Reproducibility (CoRR), and the Autism Brain Imaging Data Exchange (ABIDE) datasets and the results have been posted online.

The purpose of CoRR is to provide an open-science dataset to assess the reliability of functional and structural connectomics by defining test-retest reliability of commonly used MR metrics; to understand the variability of these metrics across sites; and to establish a standard benchmark dataset on which to evaluate new imaging metrics [17]. PCP-QAP normative data for the CoRR study was downloaded from https://github.com/preprocessed-connectomes-project/quality-assessment-protocol. The Mindcontrol database was populated with pointers to 2,963 CoRR structural images residing on an Amazon S3 bucket along with their corresponding PCP-QAP metrics. The demo of the CoRR dataset with PCP-QAP metrics is hosted at http://mindcontrol-corr.herokuapp.com.

The overarching goal of the ABIDE initiative is to expedite the discovery of the neural basis of autism by providing open access to a large, heterogeneous collection of structural and functional neuroimaging datasets collected from over 23 institutions [18]. The Preprocessed Connectomes Project provides cortical thickness measures of the ABIDE dataset output by the ANTs software package [19], along with summary statistics across regions of interests (ROIs) defined by the Desikan-Killiany-Tourville (DKT) protocol [20]. The Mindcontrol database was populated with pointer URLs to S3-hosted cortical thickness images and their corresponding ROI summary measures, along with PCP-QAP metrics. The demo of the ABIDE dataset with ANTS cortical thickness and PCP-QAP metrics is located at http://mindcontrol-abide.herokuapp.com.

## 4. Discussion

Mindcontrol is a configurable neuroinformatics dashboard that links study information and descriptive statistics with scientific data visualization, MR images, and their overlays (segmentation or otherwise). The three configurations demonstrated in this report show the link between MRI quality metrics and raw data, the link between Freesurfer regional volumes and segmentation quality, and the link between ANTS cortical thickness summary statistics and segmentation/thickness estimates of the volume. The platform is configurable, open-source, and software/pipeline agnostic, enabling researchers to configure it to their particular analyses. The dashboard allows researchers to assign editing tasks to others, who can then perform edits on the application itself.

The Mindcontrol platform streamlines and standardizes QC procedures. The traditional method of collaborative QC within a lab assembles disparate software components into a procedure that is vulnerable to clerical errors. The QC operators use existing viewers (such as FSLview or Freeview) to view and edit the images, making notes on a collaborative spreadsheet application (such as google-docs). They must carefully adhere to a common folder structure and naming convention so that other lab members and any automated processing scripts can locate these images. Distributions of scalar metrics are then plotted to identify outliers using a separate data analysis software program. The results of that analysis must then be reviewed using the imaging software to ensure that outliers are appropriately screened. This method is inherently inefficient because images must be loaded multiple times and attention split between the imaging platform and annotation software. Additionally, results of the QC must be maintained consistently across several software packages. Clerical errors are common and time-consuming to resolve because naming convention is not explicitly enforced, and manual edits could be lost within the filesystem without thorough documentation by research assistants. Google-spreadsheets are collaborative, but ideally this pass/fail/edited QC information would be directly linked to the data. Mindcontrol stores all notes, annotations, and QC results, and in-browser edits internally (Mongo database backend). User edits can be extracted automatically and written to the filesystem, eliminating the potential for clerical errors. An example python script to do this can be found at https://github.com/akeshavan/mindcontrol/wiki/Applying-Painter-Edits. Scalar metrics are linked to 3D images, enabling a user to inspect an outlier image with the click of a button. Mind-control is web-based, so it can be used on any device; QC operators can even use a tablet with stylus to edit, which is more natural than using a mouse.

There have been considerable efforts in this field to ensure data quality on a large scale. The human connectome project’s extensive informatics pipeline, which includes a database service, QC procedures, and a data visualization platform, has been key to the project’s success in collecting a large, high-quality dataset [21]. The Allen Brain Atlas offers a comprehensive genetic, neuroanatomical, and connectivity web-based data exploration portal, linking an MRI viewer to data tables [22]. The open-source LORIS web-based data management system integrates an image viewer with extensive QC modules [23]. Mindcontrol supplements these efforts by providing a lightweight and extensible data management and visualization system with the added capability to perform edits and curate annotations within the application.

Table 4 shows examples of subjects from the FCP, CoRR and ABIDE datasets with low-quality scans or segmentations, identified using Mindcontrol. The tails of various PCP-QAP metric distributions for both the ABIDE and CoRR datasets could be filtered interactively to isolate images with motion artifacts, extensive blurring, and noise. In the ABIDE dataset, filtering by the entropy focus criterion (EFC) greater than 17 identified images with extreme motion artifacts. The range of the EFC for the CoRR dataset was much smaller (less than 2) and the image with the highest EFC had no motion artifacts, but failed QC due to excessive defacing. In the ABIDE dataset, filtering for the high FWHM extremes identified images with motion artifacts, grainy/noisy images, and one extremely blurry image. On the other hand, in the CoRR dataset, the image with high FWHM had an extreme bias field. In the CoRR dataset, the images with very low contrast-to-noise (CNR) had motion artifacts, while the ABIDE images did not. Overall, examining the extremes of the PCP-QAP metrics with Mindcontrol identified outliers, but the relationship between artifacts and metrics varied by study.

Exploring the link between ANTS Cortical Thickness and the PCP-QAP metrics in the ABIDE dataset, we found that selecting the higher tail of average left-and right-postcentral gyrus thickness corresponded to datasets at the higher range of PCP-QAP FWHM. Conversely, selecting the lower tail of the precentral and postcentral gyrus thicknesses related to the lower range of the FWHM. It was difficult to pinpoint errors in the ANTS Cortical Thickness segmentation images because the data was normalized to MNI space. In the future, it would be better to QC each step of the ANTS pipeline to ensure that 1) segmentation in native space was accurate and 2) normalization to MNI space was reasonable.

Mindcontrol is particularly useful to investigate where errors occurred in segmentation algorithms. In the FCP dataset, the most common errors in segmentation with Freesurfer were that 1) parts of the temporal lobes were excluded from the segmentation and 2) the gray matter segmentation entered the dura. Scans with low-quality temporal lobe segmentation were found by selecting the lower tail of the amygdala or temporal pole volume distributions. Often, these images exhibited poor gray/white contrast in the temporal pole region, and low signal to noise. Initially, we observed that dura missclassification occurred most frequently in the precentral and postcentral gyri. We then selected data points with high precentral/postcentral volumes to locate these errors. However, scans in the middle of these metric distributions also exhibited dura misclassification, suggesting that this particular problem may be consistent across the entire dataset. In this example, it is necessary to QC every scan, regardless of where its summary metrics lie on the distribution.

In cases where every scan must be quality controlled, Mindcontrol’s summary statistic distributions and annotation features serve as a triage tool, sorting cases that are likely to require more time or expertise. When training new editors, Mindcontrol’s annotation and notes features enable users to ask questions, mark the image with the point or curve annotation feature, and assign images to more senior editors to review and provide feedback. An initial Mindcontrol quality check can be used to estimate total editing time and expertise needed for a study, enabling a strategic allocation of resources. Leveraging Mindcontrol as an integrated quality control tool can make processing methods more efficient, organized, and collaborative.

**Table 1:** A table of bad quality data or segmentations found on Mindcontrol

| Dataset | Filter | Algorithm | Example Subject IDs | Observation |
| --- | --- | --- | --- | --- |
| ABIDE | Higher end of FWHM | PCP-QAP | 50528, 0050511, 0050519 | Motion artifact |
| ABIDE | Higher end of FWHM | PCP-QAP | 50496 | Very grainy |
| ABIDE | Higher end of FWHM | PCP-QAP | 50611 | Extremely blurry |
| ABIDE | High EFC | PCP-QAP | 51160, 0051191, 0051166, 0051174, 0051192, 0051165, 0051186 | Motion artifact |
| ABIDE | High QI1 | PCP-QAP | 50197, 50017 | Very noisy, motion artifact |
| CoRR | Lower end of CNR | PCP-QAP | 25073, 25085, | Motion artifact |
| CoRR | High EFC (range much lower than ABIDE) | PCP-QAP | 25567 | No motion artifact, but frontal lobe cut off (excessive defacing) |
| CoRR | High FWHM | PCP-QAP | 27040 | Needs major bias field correction |
| FCP | Lower end of Left-Amygdala, temporal-pole, Left-Amygdala distribution | Freesurfer | sub48830, sub93262, sub55176, sub75919 | Temporal lobes not correctly segmented; gray white delineation difficult to see |
| FCP | Higher end of SuperiorFrontal, Precentral, Postcentral thickness | Freesurfer | sub98317, sub27536, sub28795, sub10582, sub93975 | Gray matter segmentation enters dura |

## 5. Future Directions

Mindcontrol is being actively developed to incorporate new features that will improve outlier detection, efficiency, and collaboration. New information visualizations to detect outliers include: scatter plots to compare two metrics against each other, and a longitudinal view of a single-subject trajectory for a given metric to detect uncharacteristic temporal changes. New scientific data visualizations are planned using the BrainBrowser library [24] to display cortical surfaces. A beta version of real-time collaborative annotations, where two users can annotate the same image and see the edits of the other user as they occur, is in the testing phase.

Currently, configuring Mindcontrol involves creating one JSON file to describe the different modules and another JSON file to populate the Mongo database with pointers to images and their scalar metrics. In the future, this process could be streamlined by creating a Mindcontrol configuration for datasets with a standardized folder structure, like the Brain Imaging Data Structure (BIDS) [25], and their BIDS-compliant derivatives [26]. Additionally, implementing the server-side application within a container, like Docker, will make it easier to deploy a Mindcontrol server. Further development of Mindcontrol will include the flexible importing of additional scalar metrics, such as measures of structural complexity, calculated by third-party toolboxes developed to complement standard analysis pipelines [27, 28]. This will enable researchers to collaborate on the same dataset by uploading metrics from their newly developed algorithms, and will enable them to easily explore their results in the context of metrics contributed by others. Finally, Mindcontrol has the potential to be a large-scale crowd-sourcing platform for segmentation editing and quality control. We hope the functionality, ease-of-use, and modularity offered by Mindcontrol will help to improve the standards used by studies relying on brain segmentation.

## 6. Software Availability

The Mindcontrol codebase is hosted on GitHub at http://github.com/akeshavan/mindcontrol, along with installation instructions. The Mindcontrol configuration of the FCP data is located on the master branch of the GitHub repository, and the configurations for CoRR and ABIDE are located at http://github.com/akeshavan/mindcontrol_configs, along with configuration documentation. Mindcontrol is licensed under the Apache License, Version 2.0.

## 7. Acknowledgements

AK would like to acknowledge Satrajit Ghosh and Arno Klein for their mentorship, the Papaya.js developers for their support, and Cameron Craddock for his assistance with the CoRR and ABIDE datasets. AK, ED, IM, KJ, and CM would like to thank the organizers of Neurohackweek and the Brainhack events for fostering a collaborative environment in which to develop new ideas and software. CM was supported by a fellowship from the Canadian Institutes of Health Research (FRN-146793). ED and KJ were supported by the National Defense Science and Engineering Graduate Fellowship. Icons from Figure 1 were sourced from https://icons8.com.

